# Biochemical and structural characterisation of a family GH5 cellulase from endosymbiont of shipworm *P. megotara*

**DOI:** 10.1101/2023.01.17.521928

**Authors:** Madan Junghare, Tamilvendan Manavalan, Lasse Fredriksen, Ingar Leiros, Bjørn Altermark, Vincent G.H. Eijsink, Gustav Vaaje-Kolstad

**Author notes:** Corresponding authors (M. Junghare), (G. Vaaje-Kolstad).

## Abstract

Cellulases play a key role in enzymatic conversion of plant cell-wall polysaccharides into simple and economically relevant sugars. The discovery of novel cellulases from exotic biological niches is of interest as they may present properties that are valuable in biorefining of lignocellulose. We have characterized a glycoside hydrolase 5 (GH5) domain of a bi-catalytic GH5-GH6 multidomain enzyme from the unusual bacterial endosymbiont *Teredinibacter waterburyi* of the wood-digesting shipworm *Psiloteredo megotara*. The cellulase enzyme, *Tw*Cel5, was produced with and without a native C-terminal family 10 carbohydrate-binding module belongs to GH5, subfamily 2. Both variants showed hydrolytic endo-activity on soluble substrates such as, β-glucan, carboxymethylcellulose and konjac glucomannan. However, low activity was observed towards a crystalline form of cellulose. Interestingly, when co-incubated with a cellulose active LPMO, a clear synergy was observed that boosted hydrolysis of crystalline cellulose. The crystal structure of the GH5 catalytic domain was solved to 1.0 Å resolution and revealed a substrate binding cleft containing a putative +3 subsite, which is uncommon in this enzyme family. The enzyme *Tw*Cel5 was active in a wide range of pH and temperatures and showed high tolerance for NaCl. This study provides an important advance on discovery new enzymes from shipworm and shed new light on biochemical and structural characterization of cellulolytic cellulase and showed boost in hydrolytic activity of cellulase on crystalline cellulose when co-incubated with cellulose active LPMO. These findings will be relevant for the development of future enzyme cocktail that may be useful for the biotechnological conversion of lignocellulose.

## Background

The desire to reduce the consumption of fossil fuels has sparked massive interest in searching alternative renewable energy sources, including lignocellulosic biomass. Plant-based lignocellulosic biomass is an abundant polymeric material that may be exploited for the production of renewable energy thus helping to reduce the consumption of fossil fuels. Lignocellulose consists of polysaccharides, including cellulose, hemicelluloses, and pectin, as well as lignin, a complex aromatic polymer. Cellulose is the most important structural component in plant biomass with an estimated global production of more than 1.5 × 10^12^ tons per year [1]. Cellulose is a homopolymer of glucose, linked by β-1,4-glycosidic bonds [2, 3]. Cellulose chains associate into insoluble, often crystalline fiber structures, which make the material strong and difficult for biodegradation. Intermolecular interactions with the other polymeric cell wall compounds add to this strength and recalcitrance. Consequently, although cellulose is an attractive source of green energy, its exploitation is intricate due to its resistance to enzymatic depolymerization [4]. Indeed, an efficient conversion of lignocellulosic biomass, or even relatively pure cellulose fibers, requires multiple enzymes acting synergistically to deconstruct the feedstock and to generate monomeric sugars that can subsequently be converted to fuels and chemicals [5].

Enzymatic cellulose depolymerization is catalysed by cellulases and lytic polysaccharide monooxygenases (LPMOs) acting in concert [5]. The cellulases include endo- and exo-acting glycosyl hydrolases (GHs) that are though to act synergistically because, for instance, endo-β-1,4-glucanases hydrolyze internal glycosidic bonds to generate new chain ends on which exo-β-1,4-glucanases, also known cellobiohydrolases, can act. Despite decades of research, there is still a need for novel and efficient cellulases that may help improving sustainability and economy of biorefining processes. Until now, most cellulose-active enzymes have been isolated and characterised from wood-decaying fungi and soil bacteria [6-9]. Cellulose-degrading higher organisms may use symbiotic microbes as a source of enzymes for biomass degradation, such as the marine shipworm which is an eminent lignocellulose degrader [10-12], remain largely unexplored. In the present study, we analysed marine wood-digesting bivalve molluscs called shipworms, which feeds on submerged wood in the ocean.

The shipworms are marine molluscs of the order *Myida* and the family *Teredinidae* (also called “termites of the sea”). They are wood-boring bivalves found throughout the world’s oceans [12]. They are notorious for boring into wooden structures immersed in sea water, where they settle on and begin to excavate into wood as larvae that grow eventually to become elongated worms [13]. Thus, shipworms are unique because only few organisms have evolved the ability to feed on woody biomass as the sole nutrient source [7]. Majority of shipworm species possess a simple digestive system with a large caecum and a short intestine. Previously it was suggested that shipworms may present few microbes in their digestive systems (cecum) that help in wood digestion [14]. However, later it was reported that endosymbiotic bacteria residing in a specialized region of the gill tissue have been shown to fix atmospheric nitrogen [15] but also produce variety of carbohydrate active enzymes (CAZymes) that function in lignocellulose deconstruction in shipworms [16]. In addition to this, shipworms themselves also produce several endogenous CAZymes that are secreted by a specialized digestive gland that finally accumulate in cecum for lignocellulose digestion [10, 11]. Because of these unique features, shipworms are an attractive target for the discovery of new CAZymes for depolymerization of lignocellulose.

The genome analysis of endosymbiotic bacteria in combination with proteome analysis from cecum of shipworms revealed that endosymbionts harbour large number of cellulases that are classified into several glycosyl hydrolase (GH) families with different substrate specificities [16-18]. For instance, GH5 which is a large protein family contains not only endo-glucanases (EC 3.2.1.4), but also β-mannanases (EC 3.2.1.78), exo-1,3-glucanases (EC 3.2.1.58), endo-1,6-glucanases (EC 3.2.1.75), xylanases (EC 3.2.1.8), endo-glycoceramidases (EC 3.2.1.123) and xanthanase [19]. In the quest for novel cellulases, we have performed the identification, cloning, expression, functional analysis and structural characterization of a cellulase belonging to glycoside hydrolase family 5 (GH5). This protein, share high homology to putative GH5 cellulase from the shipworm endosymbiont *Teredinibacter waterburyi*, named *Tw*Cel5, is part of a large multi-domain cellulase comprised of a N-terminal GH5 domain, followed by three CBM10 domains separated by serine-rich linker region and a C-terminal GH6 domain. In this study, we have produced, and functionally compared two variants of the GH5 enzyme: the catalytic domain only (*Tw*Cel5_CAT_) and the catalytic domain connected to a C-terminal CBM10 (*Tw*Cel5_CBM_). Next to functional characterization of the two variants, we also explored the potential synergistic effect between GH5 cellulase and Cels2, a cellulose-active bacterial LPMO [20], boosted activity for depolymerization of crystalline cellulose. The results showed that *Tw*Cel5 is an endo-β-1,4-glucanase with catalytic properties profile that renders it as a potentially attractive industrial biocatalyst for cellulose bioconversion.

## Results and discussion

### Sequence analysis and structural modelling

The 3312 nucleotides gene sequence (GenBank; OP793796) encode a multidomain protein (WAK85940.1), *Tw*Cel5-6, consisting of 1103 amino acids residues that possess a putative N-terminal signal peptide for protein secretion. The deduced amino acid sequence analysis using Interpro classified *Tw*Cel5-6 protein into putative N-terminal glycosyl hydrolase family 5 (GH5) domain (amino acids 15-322) and a C-terminal glycosyl hydrolase family 6 (GH6) domain (amino acids 690-1103), which are interspaced by 368 amino acids long region encoding three cellulose binding CBM10 modules (Fig. 1A). The five modules are connected by poly-serine linkers that are thought to be flexible, disorganized spacers [21].

**Fig. 1.**
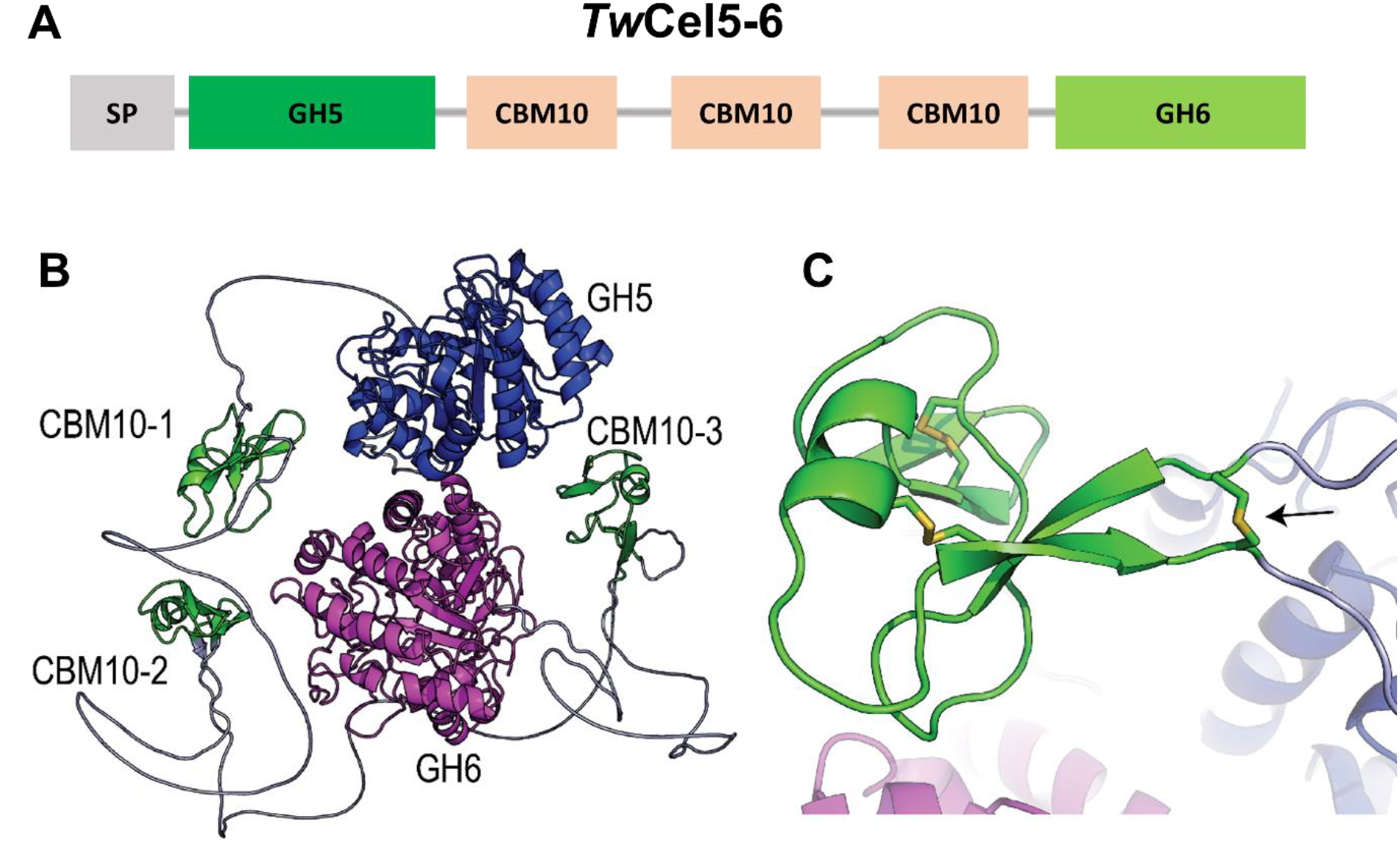
Multi-modular architecture of the *Tw*Cel5-6. (A) Display of the full-length gene encoding protein *Tw*Cel5-6 is composed of a signal peptide (SP), a glycosyl hydrolase 5 (GH5), three family 10 cellulose-binding modules (CBMs), and a glycosyl hydrolase 6 (GH6) catalytic domains, respectively. (B) Alphafold2 predicted structure of *Tw*Cel5-6; the unstructured linkers, CBM10s and the GH5 and GH6 catalytic modules are colored grey, green, blue, and magenta, respectively. (C) Details of the first CBM10 module showing the disulphide bridge (black arrow) connecting the linkers that enter and exit the module.

A BLAST search using the deduced amino acid sequence of *Tw*Cel5-6 revealed the closest 99.49 % sequence identity with a hitherto uncharacterized glycosyl hydrolase (WP_223144885.1) from the shipworm gill endosymbiont *Teredinibacter waterburyi* [22]. Notably, the latter partial protein only comprised of 773 amino acids residues, contain N-terminal putative GH5 domain (amino acids 15-322) and incomplete C-terminal GH6 domain (amino acids 688-773) which are interspaced by three cellulose binding CBM10 modules. Alphafold2 structure prediction [23] of *Tw*Cel5-6 through Colabfold [24] showed, as predicted, an unstructured N-terminus (most likely the signal peptide) followed by five modules all connected by flexible linkers (Fig. 1B; see additional file 1: Fig. S1 for prediction quality). Interestingly, Alphafold modelling predicts that the two first CBM10 modules contain a disulfide bridge that connects the end of the N-terminal linker with the start of the C-terminal linker (Fig. 1C).

### Protein expression, purification, and activity screening of TwCel5

Attempts to produce the full-length protein and several truncated variants failed (data not shown), except for the GH5 catalytic domain alone, *Tw*Cel5_CAT_, and the catalytic domain connected to one CBM10, *Tw*Cel5_CBM_, which were both expressed in soluble and active form. cDNA encoding the GH5 catalytic module of the full-length protein was cloned without the predicted signal peptide (Fig S1). Both proteins carried an affinity tag, such as *Tw*Cel5_CAT_ contained C-terminal 6xHis-tag and *Tw*Cel5_CAT_ contained N-terminal V5-6xHis-tag that were purified to homogeneity by using affinity chromatography followed by size exclusion chromatography. SDS-PAGE analysis of the purified proteins showed single homogenous protein bands migrating at around ∼ 35 kDa and ∼ 45 kDa for *Tw*Cel5_CAT_ and *Tw*Cel5_CBM_ (Fig. 2A), which is in accordance with the calculated theoretical protein masses of 33.7 kDa and 44.9 kDa, respectively.

**Fig. 2.**
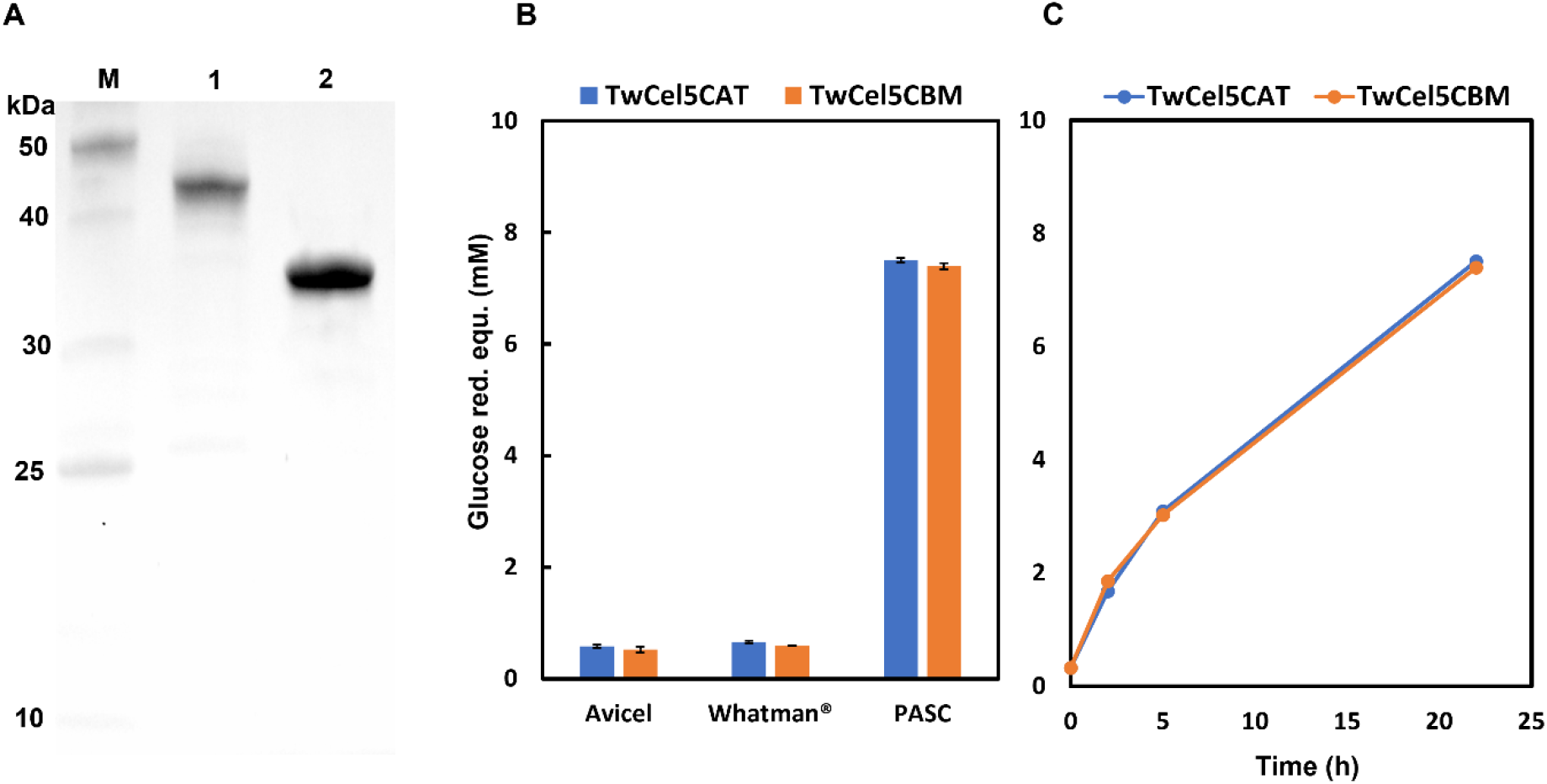
A) SDS-PAGE analysis of purified *Tw*Cel5. Lane 1, standard molecular weight markers (kDa); lane 2, *Tw*Cel5_CBM_; lane 3, *Tw*Cel5_CAT_. B). Screening of hydrolytic activity on three cellulosic 0.5 % (w/v) substrates. C). Comparative progress curves for the degradation of 0.5 % (/w/v) PASC. Reactions were performed in 50 mM sodium phosphate buffer (pH 7.5) containing 0.5 M NaCl and 1 μM enzyme incubated at 30 ºC. Hydrolytic activities were determined by measuring the release of reducing sugars using DNSA assay.

Based on CAZy database (available at http://www.cazy.org/GH5.html), the GH5 family contains a wide range of enzymes acting on diverse β-linked substrates, including cellulose. Initial activity screening experiments indicated that the two enzyme variants of *Tw*Cel5 was able to hydrolyse amorphous cellulose (PASC), but only negligible hydrolytic activity observed towards crystalline Avicel and Whatman® paper (Fig. 2B). The absence of enzymatic activity toward crystalline cellulosic substrates, indicates that the *Tw*Cel5 variants have a preference for the amorphous regions of cellulose PASC similar to, e.g., the KG35 (GH5) endo-β-1,4-glucanase that was obtained from the black-goat rumen [25]. The two variants showed comparable activities for all three substrates, which is somewhat surprising since one would expect that the presence of a CBM10 in *Tw*Cel5_CBM_ is beneficial for activity on insoluble substrates, particularly at the low substrate concentrations used in this study [26-28]. The similarity in activity was confirmed by recording comparative progress curves for PASC (Fig. 2C). This result may be taken to indication that the CBM10 has little affinity for PASC substrate than cellulose or that function of this domain cannot be observed with this truncated variant of *Tw*Cel5-6.

### Enzymatic activity on other polysaccharides

The substrate specificity of *Tw*Cel5 was assessed by measuring the hydrolytic activity on nine different soluble cellulosic and hemicellulosic substrates. Both variants showed highest activity for reactions with mixed linkage β-1,3, β-1,4 β-glucan (Fig. 3). Furthermore, the two enzyme variants also exhibited (lower) activity on CMC and konjac glucomannan that contain β-1,4 glycosidic linkages, although the enzyme without CBM produced slightly higher amounts of reducing sugars. Both enzymes were inactive on hemicellulosic substrates includes xylan, arabinogalactan, arabinan, gum arabic, xyloglucan, and lichenan (Fig. 3). It thus seems clear that the two enzyme variants prefer substrates with β-1,4-linked glucose monomers. This is in line with previous findings for several GH5 enzymes that showed similar substrate specificities

**Fig. 3.**
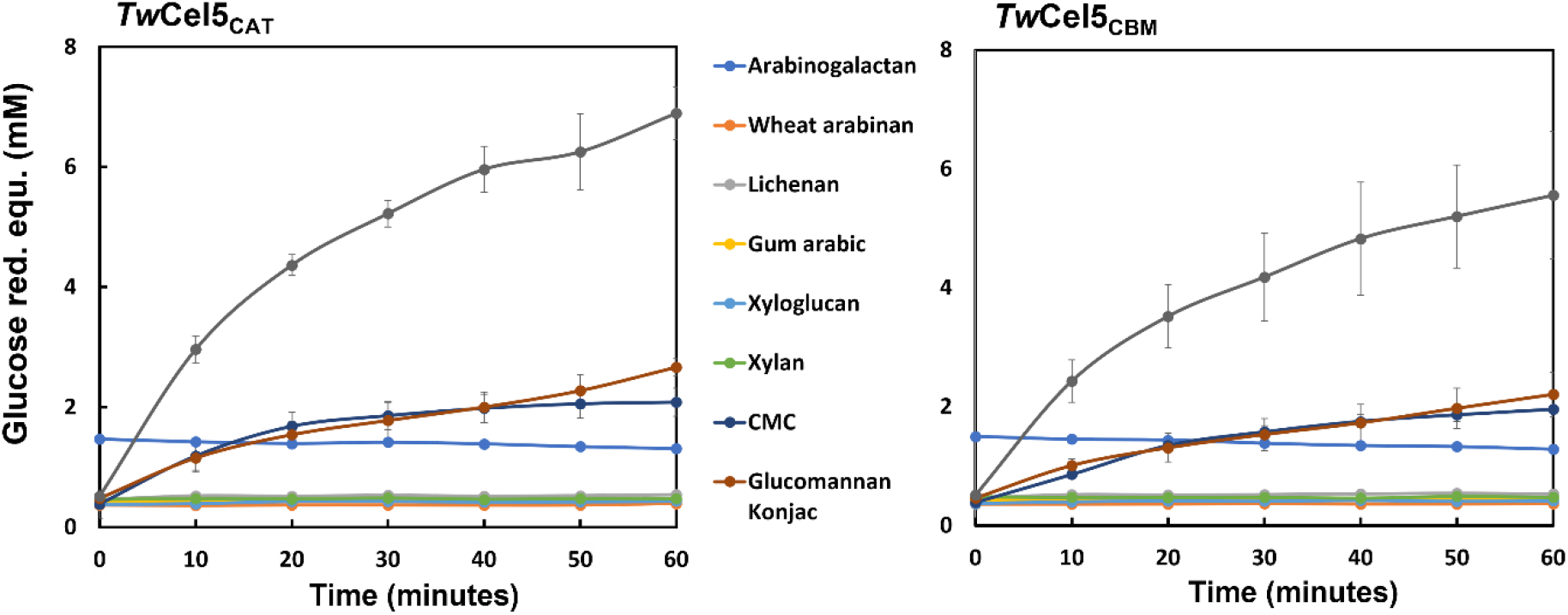
Hydrolysis of cellulosic and hemicellulose substrates by *Tw*Cel5_CAT_ and *Tw*Cel5_CBM_ as a function of time. Reactions were performed in 50 mM sodium phosphate buffer (pH 7.5) containing 0.5 M NaCl and 50 nM enzyme incubated at 30 ºC for 1 h in triplicates. Reducing sugar equivalents were quantified using glucose as a standard. The results correspond to mean and standard deviations of triplicates.

[29-31]. Of note, the enzyme showed no detectable activity toward ivory nut mannan (data not shown), which is a pure mannose polymer linked via β-1,4 glycosidic linkage. It is thus likely that β-1,4 glycosidic bonds between glucose containing substrates are the target for the *Tw*Cel5 enzyme when hydrolysing konjac glucomannan, which consists of β-1,4-linked mannose, and glucose residues in a 60:40 ratio. These biochemical properties highlight the high specificity of *Tw*Cel5 towards glucans for *P. megotara* in the context of metabolism and nutrition.

### The products analysis from enzymatic hydrolysis

To gain more insight into the mode of action of *Tw*Cel5, products generated by the enzyme after 60-minute incubation with barley β-glucan and konjac glucomannan were analysed by MALDI-TOF mass spectrometry. The MALDI-TOF MS spectra showed that *Tw*Cel5_CAT_ hydrolysed barley β-glucan to products ranging from a degree of polymerization from 4 (DP4) to DP14 (rather than only low DP products), suggesting that the enzyme mostly attacks internal glycosidic linkages (Fig. 4A, upper panel). Notably, products with DP6, DP9 and DP12 were not detected. Although this does not exactly shows how the β-glucan is hydrolysed, (barely β-glucan is a linear homopolysaccharide of consecutively linked β-(1,4)-glucosyl residues, i.e., oligomeric cellulose segments, that are separated by single β-(1,3)-linkages [32]. However, it does show that certain linkages, likely β-(1,3) linkages, are not cleaved by the enzyme. The product spectrum for konjac glucomannan showed a continuum of oligosaccharides ranging from DP4 to DP14 (Fig. 4A, lower panel), which one would expect if the ratio of glucose to mannan distribution is random, which is the case [33]. Konjac glucomannan contains about 5 - 10 % of the acetylated sugars [34], and the oligosaccharides observed at masses m/z 2189 ([DP13+Na+acetyl]+) and 2352 ([DP14+Na+acetyl]+), indeed suggest the presence of acetylation. Commensurate with these analyses, products generated from PASC was analysed by HPAEC-PAD showed a major cellobiose peak and lesser peaks for glucose, and cellotriose (Fig. 4B), as is commonly observed for cellulases. In summary, the observed activities, the distribution of products and the cleavage patterns strongly indicate that *Tw*Cel5 is a β-(1,4)-endoglucanase with a mode of action resembling that of other known GH5 enzymes [30, 35, 36].

**Fig. 4.**
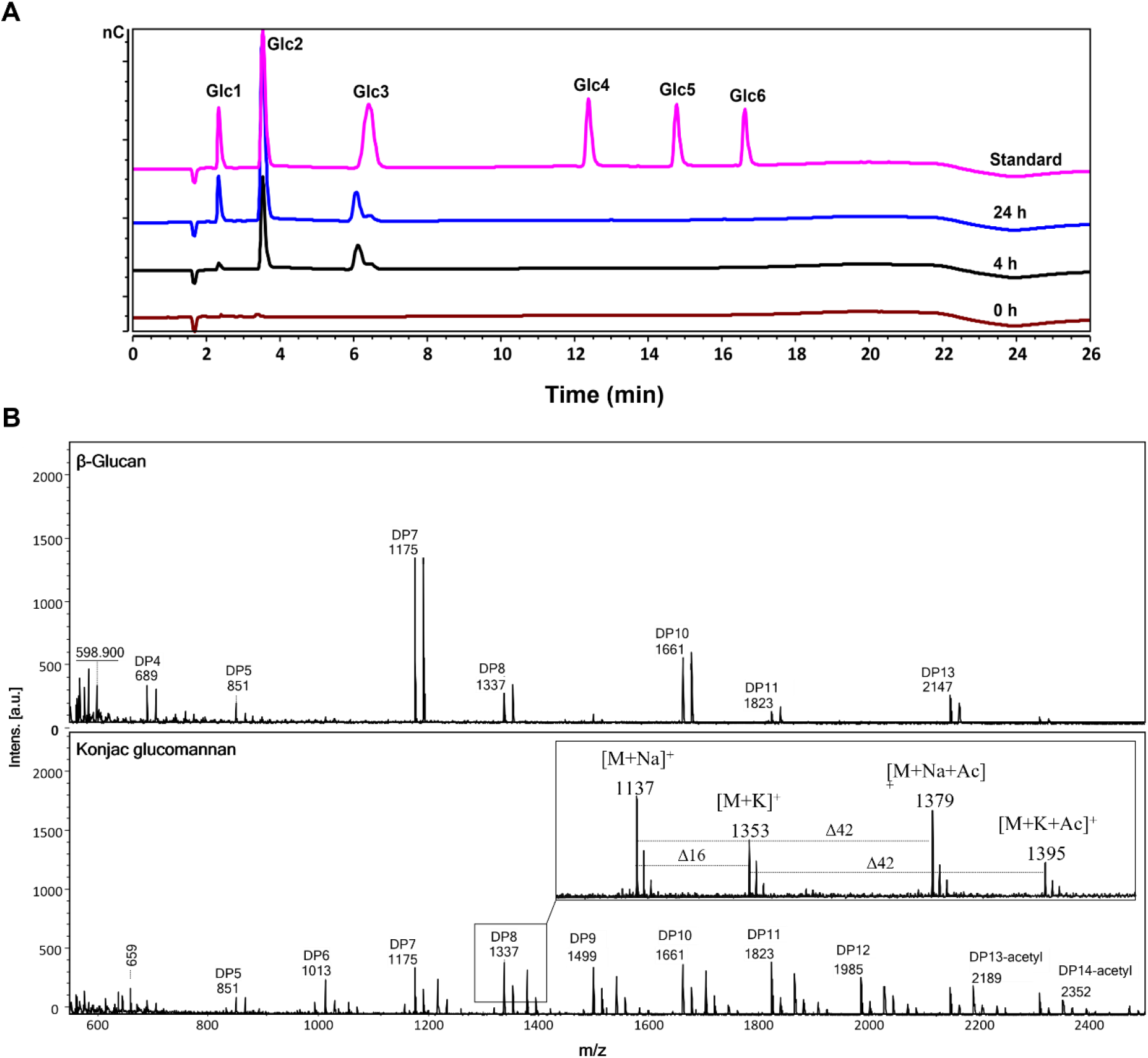
Analysis of hydrolytic products generated by *Tw*Cel5_CAT_ on different polysaccharides. A). HPAEC-PAD chromatogram showing the soluble products generated upon 0.5 % (w/v) PASC incubation of 1 μM enzyme in 50 mM sodium phosphate buffer (pH 7.5) at 30 °C for 24 h. B) MALDI-TOF MS analysis of products generated from β-glucan (upper) and konjac glucomannan (lower) 50 nM enzyme incubation in 50 mM sodium phosphate, pH 7.5 and 0.5 M NaCl for 60 min at 30 °C. The lower panel shows the Na^+^ and K^+^ adducts of oligosaccharide, and ‘DP’ stands for degree of polymerization or ‘Ac’ for acetylation. None of the labelled peaks were observed in the negative control (i.e., a reaction without enzyme). Note the *m/z* difference of 162 observed between the peaks suggest difference of a hexose residue. The products were identified based on cello-oligo standards, as shown.

### TwCel5 displays broad pH and temperature stability

Using barley β-glucan as substrate, the pH and temperature stability of *Tw*Cel5_CAT_ and *Tw*Cel5_CBM_ were determined to further characterize their optimal activities under different physicochemical conditions. When incubated at 30 ºC, both variants were active over a broad pH range, from 5 to 8, with maximum activity observed at around pH 7.0 - 7.5 (Fig. 5). More than 70 % activity was retained at between pH 4.5 and 9.6, whereas activity was drastically reduced when moving to more extreme pHs (Fig. 5). Relatively broad pH optima are commonly observed for several known GH5 enzymes, e.g., cellulase 5 from sugarcane soil metagenome [30], and endo-cellulases [31, 37], although GH5 enzymes with slightly acidic pH optima have also been reported [38, 39]. It is worth noting that the enzyme remained active for at least 60 minutes when incubated at from pH 4.0 to pH 10.6, suggesting that the protein is quite stable in a wide pH range.

**Fig. 5.**
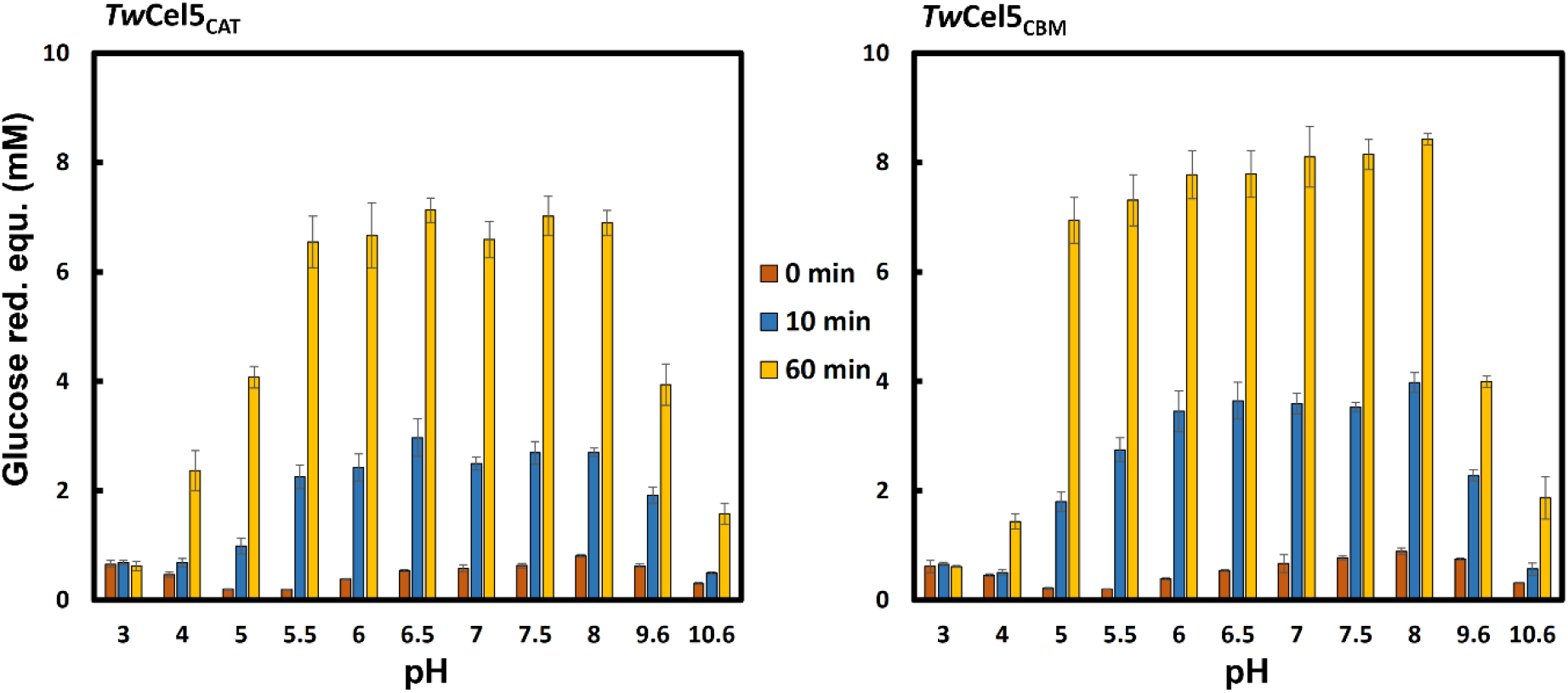
Influence of pH on the hydrolytic activity and stability of *Tw*Cel5 displaying the pH dependency of the hydrolytic activity performed in 50 mM different buffer systems using 0.5 % (w/v) β-glucan, 0.5 M NaCl and 50 nM enzyme variants, incubated at 30 ºC for 60 min. The reducing sugars values are mean and standard deviation derived from data obtained from triplicates.

Furthermore, *Tw*Cel5 activity increased in a temperature dependent manner reaching maximum activity at temperatures between 40 - 50 °C range (Fig. 6). At temperature above 50 °C signs of enzyme inactivation became noticeable during the 60 minutes incubation period and this effect became stronger for the enzyme connected with the CBM that showed slightly low activity. It is worth noting that *Tw*Cel5_CAT_ retained over 50 % hydrolytic activity at temperatures ranging from 10 °C to 60 °C, which suggests that this protein is both cold-adapted and moderately thermotolerant (Fig. 6). Although, shipworms are adapted to a cold environment, it has previously been reported that enzymes from shipworm symbionts exhibit biochemical properties similar to *Tw*Cel5_CAT_ for instance a cellulolytic endo-1,4-β-glucanase obtained from shipworm *Lyrodus pedicellatus* [37]. It is also interesting observe that, to the best of our knowledge, the combination of being active at rather alkaline pH and being thermo-tolerant is rather rare and unusual enzyme feature [40-42]. As *Tw*Cel5 is originating from a marine shipworm symbiont that reside in ocean water, we assessed the impact of NaCl salt concentration on enzyme hydrolytic activity. Using standard assay conditions, we found that the no impact of 0 - 1.5 M NaCl concentration on enzyme activity (additional file 1: Fig. S2). In conclusion, *Tw*Cel5 is a salt-tolerant β-1,4-endo-glucanase capable of function in a wide pH and temperatures ranges, which makes it an interesting candidate relevant for industrial applications.

**Fig. 6.**
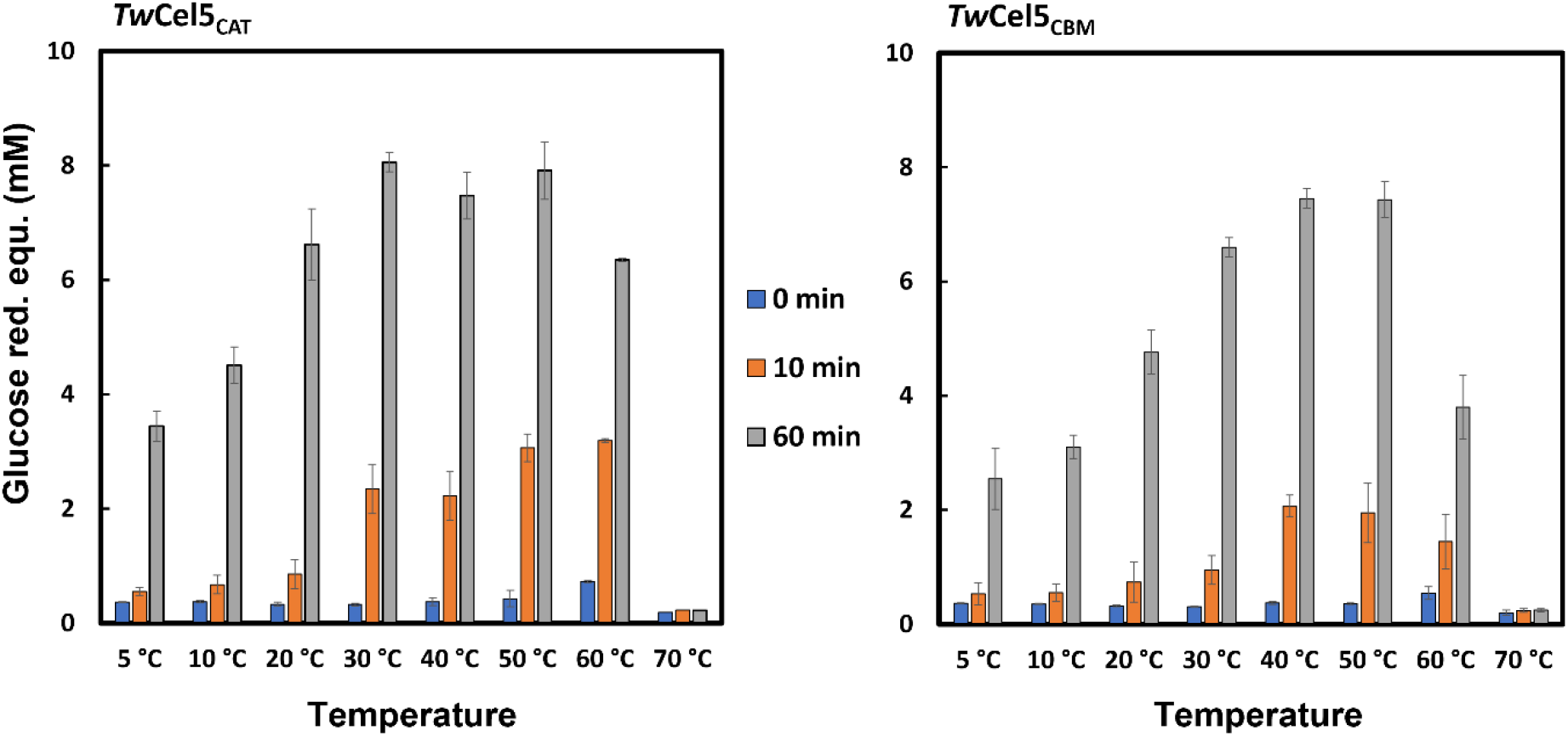
Influence of temperatures on the hydrolytic activity and stability of *Tw*Cel5 displaying the temperature dependency on the hydrolysis of β-glucan performed in 50 mM sodium phosphate buffer, pH 7.5, using 0.5 % (w/v) β-glucan, 0.5 M NaCl and 50 nM enzyme variants, incubated at different temperatures ranging from 5 ºC - 30 ºC for 60 min. The reducing sugars values are mean and standard deviation derived from triplicates.

### Co-incubation of TwCel5 with LMPO boosted hydrolytic activity

It is well known that shipworm gill endosymbionts produce a multitude of carbohydrate active enzymes, including cellulose active lytic polysaccharide monooxygenases (LPMOs) and diverse glycosyl hydrolases (GHs) to accomplish the efficient wood digestion for nutrition and growth [16, 43]. Recent study using the combination of meta-transcriptome, proteomic and biochemical analysis from wood-degrading shipworm *Lyrodus pedicellatus* reported the expression of gene encoding auxiliary activity family 10 (AA10) LPMO which likely synergise with endogenous as well as endosymbiont multi-domain glycoside hydrolases that functions in the hydrolysis of β-1,4-glucans [10]. It was thus of great interest to determine whether a typical cellulose-active AA10 LPMO, CelS2 from *Streptomyces coelicolor* A3(2) [20, 44] would enhance the hydrolytic activity of *Tw*Cel5_CAT_ or *Tw*Cel5_CBM_ specially on Avicel, a semi-crystalline form of cellulose. As expected, clear synergistic effect was observed on the generation of reducing sugars when CelS2 and *Tw*Cel5_CAT_ or *Tw*Cel5_CBM_ were combined as enzyme cocktail (Fig. 7A).

**Figure 7.**
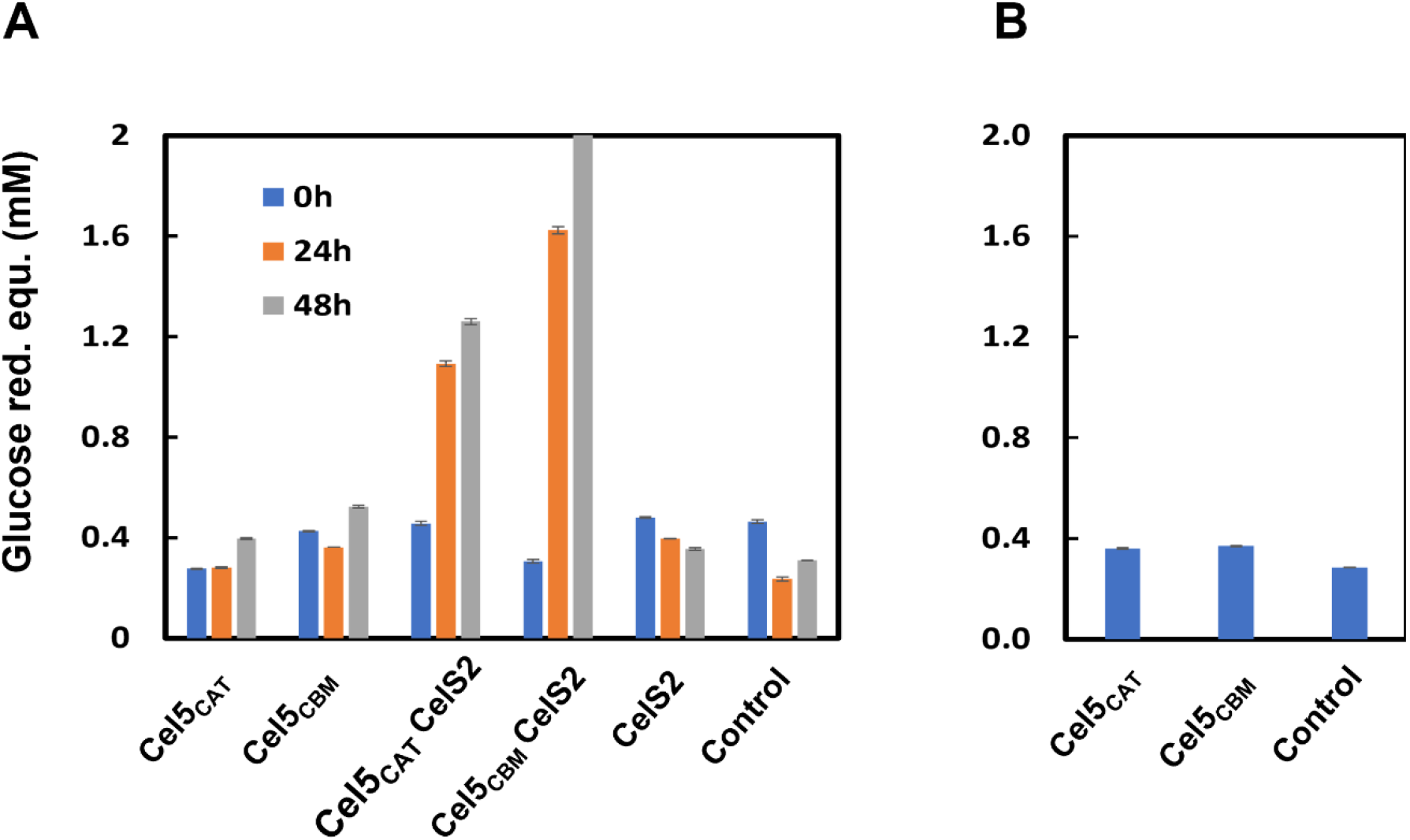
Synergy experiment between cellulase active LPMO and cellulase *Tw*Cel5 showing the boost in hydrolytic activity against Avicel. A) *Tw*Cel5_CAT_, *Tw*Cel5_CBM_ or CelS2 each 1 μM were incubated alone or in combination for different time durations, all with 1 mM ascorbate included in the reaction mixture. B) Reactions conducted from the oxidised products of avicel by CelS2 after 48 h treatment were incubated with the *Tw*Cel5 variants for 18 h; “control” refers to a reaction in which no enzyme was added. Values obtained are means and standard deviation derived from triplicate.

Interestingly, in the reaction with CelS2, the amount of reducing sugars released was higher for enzyme that connected to CBM10, *Tw*Cel5_CBM_ compared to catalytic *Tw*Cel5_CAT_. This may be taken to suggestion that LPMO activity uncovers the regions of crystalline substrate where the CBM10 is partially beneficial for the efficiency of GH5. Of note, a control reaction (Fig. 7B) showed that, potentially apparent, synergy is not just a result of GH5-catalyzed hydrolysis of longer soluble cello-oligomers generated by the LPMO. Our results add to studies showing that cellulose active LPMO boosts the activity of shipworm glycoside hydrolases on crystalline cellulose such as Avicel [43] and support the notion that LPMO action is important for wood depolymerization in shipworms for digestion and nutrition.

### Structural analysis of cellulase TwCel5_CAT_

To gain structural insights for *Tw*Cel5_CAT_, bond cleavage pattern and mode of action, we solved and determined the tertiary structure of catalytic module. *Tw*Cel5_CAT_ was crystallized in an apo form and a dataset diffracting to 1.0 Å resolution was collected (Table 1). The structure was determined by molecular replacement using the protein coordinates from the structure of the GH5 cellulase Cel5 (PDB entry 1EGZ) from *Erwinia chrysanthemi*, a gram-negative plant pathogen [45] as the search model (Table 2). *Tw*Cel5_CAT_ model was refined to R_work_ and R_free_ of 11.42 and 13.30, respectively, and the final model was deposited in the PDB database (PDB identifier 8C10). Structural comparisons using the DALI server [46] revealed highest closest structural matches to the cellulase CelE1 belonging to GH5 family (PDB identifier 4M1R; 67 % sequence identity) obtained from a sugarcane soil metagenome [30], and endoglucanase EGZ (PDB identifier 1EGZ; 67 % sequence identity) from *Erwinia chrysanthemi* [45, 47], respectively.

**Table 1:**
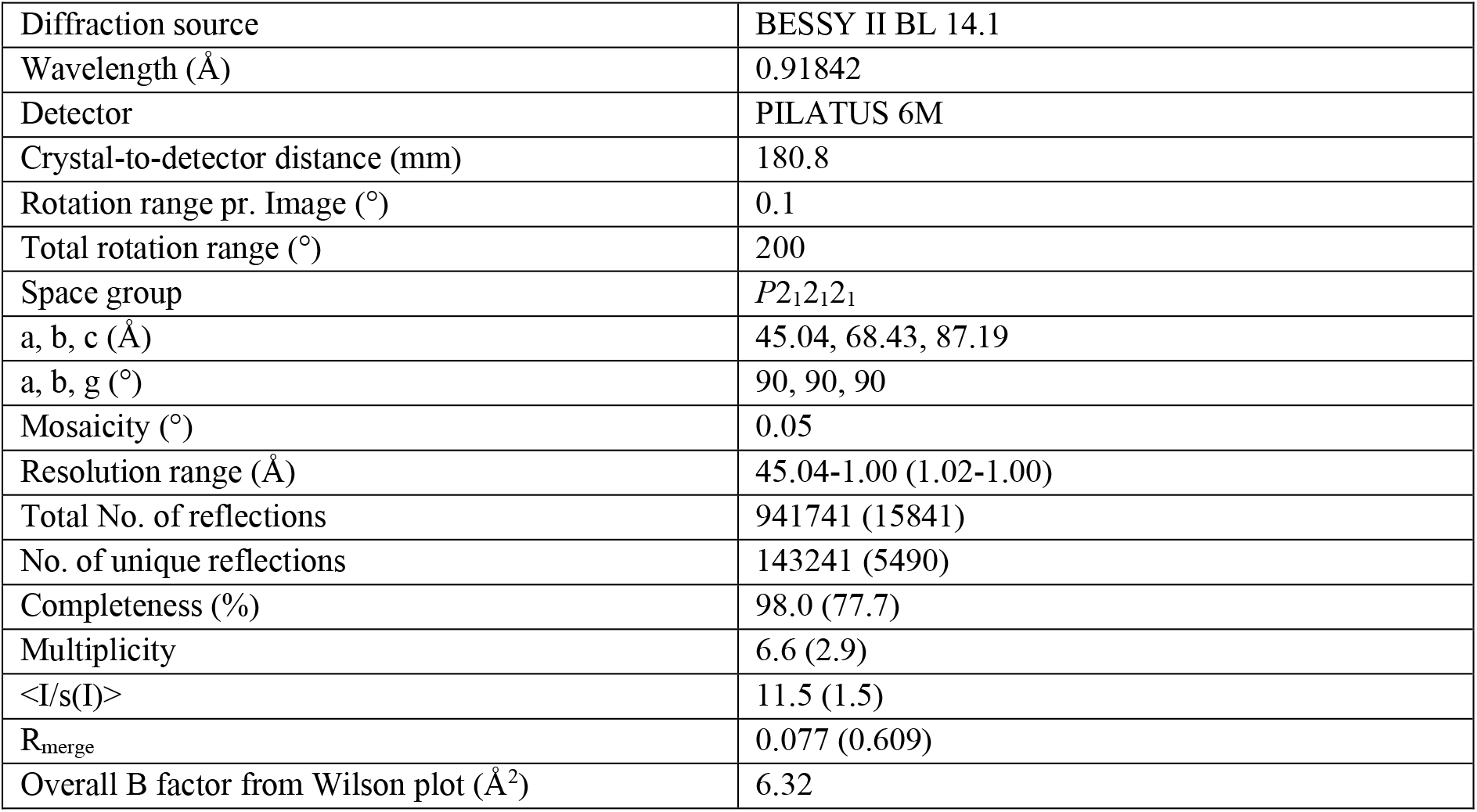
Data collection and processing statistics. Values in parentheses are for the outermost shell.

**Table 2:** Structure determination and refinement statistics. Values in parentheses are for the outermost shell.

|  |  |
| --- | --- |
| PDB code | 8C10 |
| Resolution range (Å) | 45.04-1.00 (1.02-1.00) |
| Completeness (%) | 97.85 (79.7) |
| s cutoff | None |
| No. of reflection, working set | 140934 |
| No. of reflection, reference set | 2221 |
| Final R<sub>work</sub> | 11.42 |
| Final R<sub>free</sub> | 13.39 |
| Cruickshank DPI | 0.0178 |
| No. of non-H atoms |  |
| Protein | 2427 |
| Water | 339 |
| Other* | 19 |
| Total | 2785 |
| R.m.s. deviations |  |
| Bonds (Å) | 0.0291 |
| Angles (°) | 2.31 |
| Average B factors (Å<sup>2</sup>) |  |
| Overall | 11.37 |
| Protein | 9.71 |
| Water | 22.75 |
| Other* | 20.70 |
| Ramachandran plot (%) |  |
| Preferred | 96.61 |
| Allowed | 3.39 |
| Outliers | 0 |

The first three-dimensional structure of a CAZY family GH5 subfamily 2 was a cellulase isolated from an alkaline *Bacillus* sp., found in soda lakes [48]. Consistent with the defining family member, *Tw*Cel5_CAT_ exhibits a classical (β/α)_8_-barrel fold (also known as TIM-barrel; Fig. 8A) with two conserved glutamates; Glu157 & Glu245 in the catalytic center (Fig. 8B) that are positioned to promote the expected displacement mechanism characteristic of family 5 cellulases [49]. Thus, from homology to other GH family 5 cellulases, the proton donor is expected to be Glu157, and the nucleophile is Glu228. Furthermore, substrate binding cleft of *Tw*Cel5_CAT_ is homologous to other clade 2 GH5 enzymes, showing conserved aromatic and polar amino acids involved in substrate binding (Fig. 8C). In contrast to most other GH5s subfamily 2, *Tw*Cel5_CAT_ has a tryptophan (Trp226) in the outer region of the reducing end subsites (Fig. 8D). Of the 15 unique structures from GH5s subfamily 2, only three have a Trp in this position, namely the thermostable GsCelA from *Geobacillus* sp. 70PC53 [50], the halotolerant Cel5R cloned from a soil metagenome [51] and Cel5B from *C. hutchinsonii* [52], respectively. Of note, comparison of the Alphafold2 model of *Tw*Cel5_CAT_ with the X-ray crystallographic structure showed RMSD for Ca-carbons of only 0.35 Å (Fig. S3A). Furthermore, the side chains of the amino acids in the conserved active site and substrate binding cleft were modelled clos to perfectly (Fig S3B).

**Fig. 8.**
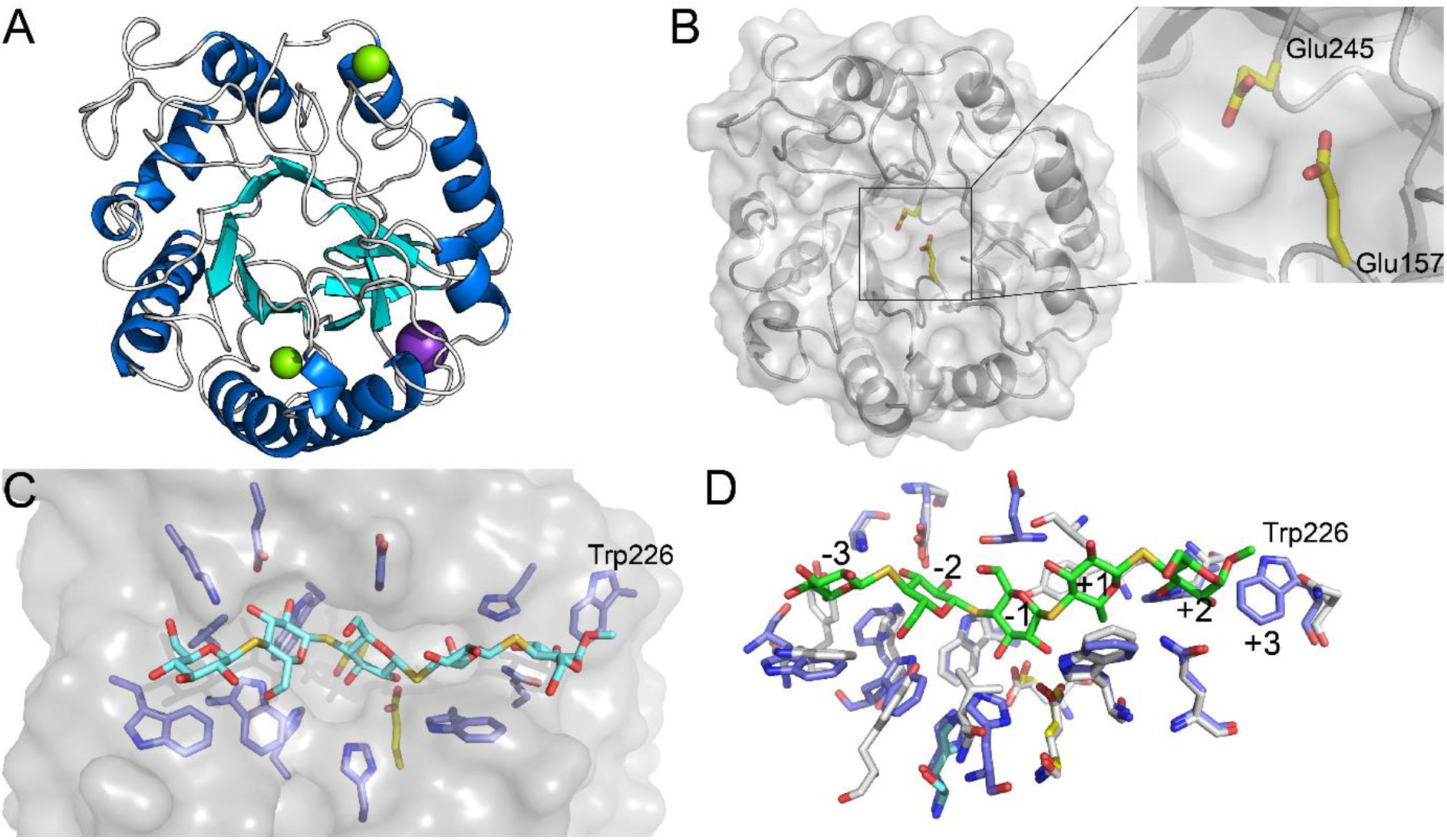
Crystal structure analysis of *Tw*Cel5_CAT_. (A). Orthogonal view of the *Tw*Cel5_CAT_ tertiary structure showing the characteristic (β/a)_8_-barrel fold. Helices are blue, β-strands cyan, and loops white are shown. Mg^++^ and K^+^ ions are shown as green- and purple-colored spheres, respectively. (B) The putative active site of *Tw*Cel5_CAT_, with the side chains of Glu157 and Glu245 shown with yellow carbons. (C) Illustration of substrate binding groove with the side chains of amino acids potentially interacting with substrate shown with blue carbons. For illustration purposes, a thiocellopentaose molecule has been placed in substrate binding site by structural superposition of *Tw*Cel5_CAT_ structure with the structure of ligand containing Cel5A from *Bacillus agaradharens* (PDB identifier: 1H5V). (D) Structural superposition of subsite defining amino acids of *Tw*Cel5_CAT_ and B. *agaradharens* Cel5A, highlighting the putative +3 subsite in *Tw*Cel5_CAT_. *Tw*Cel5_CAT_, blue carbons; Cel5A, grey carbons; substrate, green carbons.

## Conclusions

This study describes the detailed biochemical characterization and structural properties of GH5 domain of multi-domain cellulase *Tw*Cel5-6 from the shipworm’s endosymbiont *T. waterburyi* residing in the gills of shipworm including *P. megotara*. It is an endo-glucanase enzyme showing activity at broader pH from 5 - 8 and temperatures 40 - 50 °C, respectively. *Tw*Cel5_CAT_ hydrolytic activity on crystalline Avicel was boosted upon synergistic interaction with cellulose oxidizing CelS2, and in this reaction set-up the presence of a C-terminal CBM10 domain fused to *Tw*Cel5_CAT_ promoted enhanced cellulose saccharification. This endo-glucanase may be a suitable biocatalyst for liberating reducers sugars from pre-treated biomass in a broader pH and temperature range, including alkaline pH, low temperature and high salt tolerance. In summary, our study demonstrates that wood-digesting shipworms are good source of novel enzymes active at alkaline pHs and moderately thermostable cellulases.

## Methods

### Chemicals and substrates

Analytical grade substrates were used. The pre-packed 5 mL HisTrap™ affinity (HP), PD-10 desalting columns (Sephadex G-25 resin), and size exclusion chromatography column (HiLoad® Superdex, 75 pg) used for protein purification were obtained from GE Healthcare. Amorphous, phosphoric acid swollen cellulose (PASC) was prepared from Avicel according to method described before [53]. High-purity substrates include barley β-glucan, lichenan, konjac glucomannan, wheat arabinoxylan, birchwood xylan, and tamarind xyloglucan purchased from Megazyme. The model crystalline cellulose substrate Avicel PH-101, carboxymethyl cellulose (CMC), gum arabic and standard cello-oligomers purchased from Sigma-Aldrich.

### Sample collection and identification of genes

Specimens of adult shipworm *Psiloteredo megotara* collected from Norway spruce (*Picea abies*) wooden panels submerged were for about 8-9 months in the Arctic Sea near Tromsø, Norway (N 69°46’47.515”; E 18°23’53.143”). The sampling was done accordance with the Norwegian Marine Resource Act. The shipworm specimen was initially rinsed with sterile water and dissected on a clean bench to separate the specialised gill tissue containing endosymbionts. Bacterial enrichment performed using crushed gill tissues in a medium supplemented with cellulose as a carbon source that were incubated for several months. DNA was isolated from the enrichment culture using the DNeasy Blood & Tissue Kit (QIAGEN). The metagenome sequencing was carried using Illumina MiSeq 300 paired-end chemistry at the Norwegian Sequencing Centre (www.sequencing.uio.no). Analysis of contigs were assembled, annotated and uploaded to the GenBank sequence database (accession number grp 8783669). Full details of the metagenomic dataset will be published elsewhere. Genes coding for carbohydrate-active enzymes were mined using the dbCAN meta server [54]. A gene with accession number OP793796 (3312 bp) that located on BankIt2638814 Contig82 was chosen for further studies due to its novel multi-domain architecture.

### Gene cloning, expression, and protein purification

A gene (OP793796) encoding multidomain protein *Tw*Cel5-6 has an accession number WAK85940.1 (codon optimized for *E. coli* expression using the OptimumGene™ PSO algorithm) was synthesized by GenScript Biotech (Piscataway, NJ 08854, USA). Gene fragments encoding *Tw*Cel5_CAT_ and *Tw*Cel5_CBM_ were generated using through PCR using Q5 DNA polymerase (New England Biolabs, Ipswich, MA) and the primers described in Table S1. Both the genes were amplified excluding a putative signal peptide (additional file: Fig. S4), as predicted using the SignalP-5.0 prediction tool [55]. PCR products were purified using a PCR clean-up kit as per manufacturer’s instructions (Macherey-Nagel, Germany) followed by agarose (1 w/v %) gel electrophoresis. Prior to cloning, the DNA concentration was determined using a nano UV spectrophotometer (Thermo Scientific, San Jose, CA, USA). The DNA fragment encoding *Tw*Cel5_CAT_ was cloned (26 - 320 amino acid residues) into the pNIC-CH expression vector (AddGene, Cambridge, MA), which adds a C-terminal polyhistidine-tag (6xHis-tag) to the protein as per manufactures instructions. Similarly, DNA fragment encoding *Tw*Cel5_CBM_ was cloned (30 - 412 amino acid residues) into the directional champion pET151/D-TOPO™ expression vector (Invitrogen, Carlsbad, CA), which adds a cleavable N-terminal V5-6xHis-tag to the protein as per manufactures instructions.

The recombinant vectors were transformed into chemically competent OneShot™ *E. coli* TOP10 (Invitrogen, Carlsbad, CA) cells using a heat shock transformation method. Cells were grown in SOC medium for 60 minutes prior to plating on lysogenic broth (LB) agar plates supplemented with antibiotics; 50 μg/mL kanamycin (for *Tw*Cel5_CAT_) and 100 mg/mL ampicillin (for *Tw*Cel5_CBM_) depending on the vector, followed by overnight incubation at 37 °C. Colonies on the LB plates were picked and screened by colony PCR using the pair of T7 primers: T7, 5’
s-TAATACGACTCACTATAGGG-3’ and T7 reverse 5’-TAGTTATTGCTCAGCGGTGG-3’. Positive clones were picked and inoculated in liquid LB containing appropriate antibiotics (as mentioned above) and the cultures were incubated overnight at 37 °C with shaking at 200 rpm. The recombinant plasmids were isolated from the *E. coli* cells using the Zymo MiniPrep Kit (Zymo Research). Prior to transformation to expression cells, the correct integration and DNA sequence of the genes was confirmed using Sanger sequencing (GATC Biotech, Constance, Germany). The expression vectors were transformed into chemically competent OneShot BL-21 Star™ (DE3) and ArcticExpress (DE3) *E. coli* expression cells, for *Tw*Cel5_CAT_ and *Tw*Cel5_CBM_, respectively, using heat shock method as described above.

To produce recombinant proteins, *E. coli* transformants were inoculated and grown in terrific broth medium supplemented with appropriate antibiotics and cells were incubated at 37 °C with horizontal shaking (200 rpm) until the optical density (OD_600nm_) reached between 0.6 to 0.8 followed by induction by adding 0.3 mM isopropyl-β-D-thiogalactopyranoside (IPTG) and 24 h incubation at 15 °C with horizontal shaking (200 rpm). Cultures were harvested by centrifugation at 10,000 x g for 15 min at 8 °C, using a Beckman coulter centrifuge (Brea, CA, USA). Cells were stored frozen at -20 °C until further use. For protein purification, a cell-free extract was prepared by re-suspending about 5 g wet cell biomass in 50 mL of 50 mM Tris-HCl buffer, pH 7.4, supplemented with 200 mM NaCl, 10 % glycerol and 30 mM imidazole (lysis buffer). Prior to cell disruption, the suspension was supplied with 0.5 - 1 mg of cOmplete™, EDTA-free protease inhibitor cocktail (Roche) and lysozyme (0.5 mg/mL). Cells were disrupted using a cooled high-pressure homogenizer (LM20 Microfluidizer, Microfluidics). Cell debris was removed by centrifugation at 27,000 x g for 30 min at 4 °C. The resulting cell-free extracts, containing cytosolic soluble proteins, were filtered using a sterile 0.45 μm filter (Sarstedt, Nümbrecht, Germany).

The filtered cell-free extract was subjected to immobilized metal affinity chromatography using an Äkta pure chromatography system equipped with a 5-mL HisTrap HP column (GE HealthCare) equilibrated with lysis buffer (see above). After sample loading, the HisTrap column was washed extensively using 50 mM Tris-HCl, pH 7.4, supplemented with 200 mM NaCl, 10 % glycerol containing 70 mM imidazole (wash buffer) until UV absorbance dropped and stabilised at baseline level. Bound proteins were eluted using the same buffer supplemented with 500 mM imidazole (elution buffer). Eluted proteins were first analysed using SDS-polyacrylamide gel electrophoresis (SDS-PAGE) using TGX Stain-Free precast gels (Bio-Rad, Hercules, Ca, USA). The molecular weight of the recombinant proteins was estimated using Invitrogen™ BenchMark™ Pre-stained Protein Ladder (Fisher Scientific, Waltham, Massachusetts, USA). Eluted proteins were concentrated using Vivaspin® 10,000 MWCO centrifugal filter units (Sartorius, Göttingen, Germany). Proteins were purified to homogeneity using a size-exclusion chromatography column (HiLoad® Superdex, 75 pg) pre equilibrated with 50 mM sodium phosphate buffer (pH 7.4) containing 150 mM NaCl. Finally, buffer exchange was performed to 50 mM sodium citrate (pH 5.6) using a PD-10 desalting column. The protein concentration was determined by UV absorbance at 280 nm (A_280_) using theoretical molar extinction coefficients (*Tw*Cel5_CAT_: 71390 M^-1^·cm^-1^ and *Tw*Cel5_CBM_: 96745 M^-1^·cm^-1^) estimated using the ProtParam tool [56]. Purified enzymes were stored at 4 °C.

### Biochemical characterization of recombinant TwCel5

The standard assays were performed in 300 μL reaction volume containing 50 mM sodium phosphate buffer (pH 7.5), 0.5 M NaCl, 0.5 % (w/v) β-glucan and 50 nM of the purified enzyme. The reaction was started either by the addition of enzyme or substrate followed by incubation at 30 ºC using a thermomixer with horizontal agitation (500 rpm; Eppendorf, Hamburg, Germany) for 60 minutes (unless stated otherwise). To determine the pH stability and pH optima, reactions were carried out in 50 mM buffer systems (sodium citrate, pH 3.0 - 6.0; potassium phosphate, pH 6.5 - 8.0; and glycine-NaOH, 9.6 - 10.6) containing 0.5 M NaCl, 0.5 % (w/v) β-glucan and 50 nM of the purified enzyme. To determine the thermal stability and optimal temperature, reactions were performed in 50 mM sodium phosphate buffer (pH 7.5), 0.5 M NaCl, 0.5 % (w/v) β-glucan and 50 nM of the purified enzyme at various temperatures ranging from 5 - 70 °C. The effect of the NaCl concentration on activity was determined using 50 mM sodium phosphate buffer (pH 7.5), 0.5 % (w/v) β-glucan, 50 nM of the purified enzyme and various concentrations of NaCl ranging from 0 to 1.5 M. Aliquots were collected at different time intervals in a period of 60 minutes; reactions were stopped by mixing the samples immediately with DNSA reagent. Product formation was determined by quantifying the amount of reducing end sugars using the 3,5-dinitrosalicylic acid (DNSA) assay method [57] using glucose as a standard. The absorbance (A_540nm_) was recorded using Varioskan™ LUX multimode microplate reader (Thermo Scientific, San Jose, CA, USA). All the assays were performed in triplicate.

### Lignocellulose substrate specificity

The substrate specificity of purified *Tw*Cel5_CAT_ and *Tw*Cel5_CBM_ was evaluated using a wide variety of complex lignocellulosic substrates, both soluble and crystalline. The insoluble and model crystalline polysaccharides include Avicel PH-101, Whatman® cellulose filter paper (0.5 μm particle size) and phosphoric-acid swollen cellulose (PASC) was used as amorphous substrate. The soluble lignocellulosic substrates included β-glucan, birchwood xylan, carboxymethyl cellulose (CMC), wheat arabinoxylan, konjac glucomannan, xyloglucan, and lichenan. Soluble substrates were dissolved according to the supplier’s protocol. The standard reactions were carried out in 300 μL reaction volume using 50 mM sodium phosphate buffer, pH 7.5, 0.5 M NaCl, using 50 nM of enzyme for soluble substrates (0.5 % w/v) whereas 1 μM for insoluble substrates (1 % w/v) that were incubated at 30 °C for 60 minutes, for soluble substrates, or 24 hours, for insoluble substrates, with horizontal agitation (500 rpm). Aliquots were taken at different intervals; reactions were stopped by mixing the samples immediately with DSNA reagent. Release of reducing end sugars was measured using DNSA assay, using glucose as a standard, as described (see above). When using insoluble substrates samples were filtered before measurement.

### Cellulase-LPMO synergy experiment

Purified CelS2 from *Streptomyces coelicolor* was a kind gift from Dr. Zarah Forsberg [20]. The cellulase-LPMO synergy was assessed by performing reactions with crystalline Avicel (1 % w/v) in 50 mM sodium phosphate buffer (pH 6.0) using a thermomixer (Eppendorf, Hamburg, Germany) incubated at 30 °C with horizontal agitation (1000 rpm). Experiments to determine the synergy were conducted at a fixed total enzyme concentration of, such as 1 μM copper saturated CelS2 and/or 1 μM of one of the *Tw*Cel5 cellulase variants. The reactions were started by firstly supplying 1 mM ascorbic acid (final concentration) to all reaction mixtures, immediately followed by addition of the enzyme. Reactions were incubated for 48 hours, and aliquots were taken at different time intervals followed by filtration using 0.45 μm filter for removing insoluble substrate and to stop the reaction. To check for generation of reducing ends due to the action of the cellulase variants of oligomeric products solubilized by the LPMO, which could lead to a false impression of synergy, control reactions were performed in which Avicel was first incubated with the LPMO for 48 hours, after which the products were treated with the *Tw*Cel5 variants. Cellulose saccharification was assessed using the reducing end assay described above and all the experiments were performed in triplicate.

### Product analysis by HPAEC-PAD (ICS-6000)

Hydrolytic products generated from PASC were detected by a Dionex ICS6000 system (Thermo Scientific, San Jose, CA, USA) using high performance anion exchange chromatography connected to pulsed amperometric detector using CarboPac™ PA200 IC analytical column. The eluent B (0.1 M NaOH and 1 M sodium acetate) and eluent A (0.1 M NaOH) was applied using following gradient program: 0 - 5.5 % B for 3 min, 5.5 - 15 % B for 6 min, 15 - 100 % B for 11 min, 100 - 0 % B for 6 s, 0 % B for 6 min. The eluent flow rate was set to 0.5 mL/min. The cello-oligosaccharide with a degree of polymerization from one to five (DP1 - DP5), was used as standards for product identification. The data was analysed using Chromeleon 7.2.9 software.

### Product analysis by MALDI-TOF MS

Hydrolytic products generated from β-glucan and konjac glucomannan were identified using a UltrafleXtreme matrix assisted laser desorption ionization time of flight mass spectrometer (Bruker Daltonics GmbH, Bremen, Germany) equipped with a Nitrogen 337-nm laser. Samples were prepared by mixing one microliter of sample with two microliter of 2,5-dihydrooxybenzoic acid (DHB) solution (9 mg/mL) that directly mounted onto a MTP 384 ground steel target plate (Bruker Daltonics). Sample spots were allowed to dry on the plate using a flow of dry hot air. Data was acquired using Bruker flexControl and flexAnalysis software. Products were identified based on *m/z* values.

### Crystallization, data collection and analysis

Crystallization experiments were performed with a stock solution of the purified protein at 12.4 mg/mL, as estimated by A_280_, in 6 mM NaCl, 20 mM Tris-HCl at pH 8.0. Initial crystallization experiments were performed using the vapour diffusion sitting drop method set up by a Phoenix crystallization robot (Art Robbins Instruments). The crystallisation experiments were set up with sixty μl reservoir solutions and sitting drops with equal amounts of reservoir solution mixed with protein stock solution in a total drop volume of one microliter. The plates were incubated at 20 °C. Crystals appeared in 1-2 weeks in conditions containing 0.2 M MgCl_2_, 0.1 M Tris-HCl, pH 8.5, and 25 % w/v of polyethylene glycol 3350 (PEG 3350). Crystals were harvested, transferred to a cryoprotectant solution consisting of the reservoir solution supplemented with 15 % ethylene glycol and flash cooled in liquid N_2_. X-ray diffraction data were collected at BEAMLINE 14.1 at BESSY II (Berlin, Germany). Data collection and processing statistics are presented in Table 1, and structure determination and refinement statistics are presented in Table 2. The crystal structure was solved by molecular replacement using MolRep in the CCP4 program package [58] with 1egz.pdb as search model [45]. The initial refinement was executed in Refmac [59] followed by automated model improvement in Buccaneer [60]. The manual model building was done in Coot [61] interspersed by cycles of refinement in Refmac and resulted in a final R_work_/R_free_ of 11.42/13.39. The atomic coordinates and structural details have been deposited in the Protein Data Bank with the accession code 8C10. Figures presented in the results section were generated using Pymol v4.60 (www.pymol.org).

## Supporting information

file:///C:/Users/madenj/OneDrive%20-%20Norwegian%20University%20of%20Life%20Sciences/GH5_cellulase%20manuscript/BITE%20submission/Additional%20file%20

## Declarations

### Ethics approval and consent to participate

Not applicable Competing interests: Not applicable

## Acknowledgements

We gratefully acknowledge the financial support received from a grant by ERA-NET MarineBiotech, provided by the Research Council of Norway (grant number 283647), and by the NorZymeD and OXYMOD projects financed by Research Council of Norway (grant numbers: 221568 and 269408), respectively.

