## Supplementary material for "Biochemical and structural characterisation of a family GH5 cellulase from endosymbiont of shipworm *P. megotara*": file:///C:/Users/madenj/OneDrive%20-%20Norwegian%20University%20of%20Life%20Sciences/GH5_cellulase%20manuscript/BITE%20submission/Additional%20file%20

Additional file 1: Figures S1 to S4 and Table S1

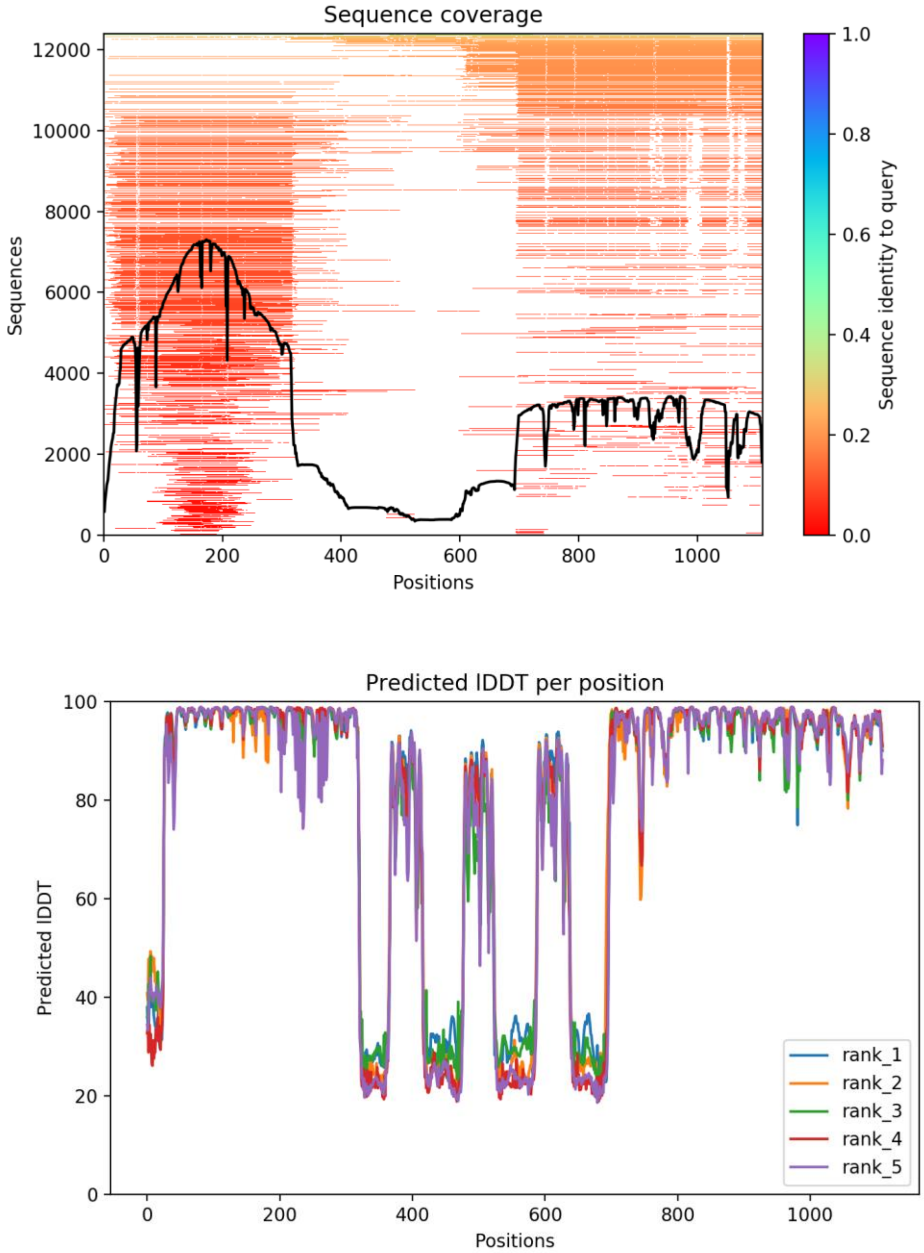


**Fig. S1.** *Tw*GH5-6 model sequence coverage and predicted LDDT. The upper panel shows the sequence coverage for the full-length protein sequence (GH5: amino acids 15-322, linkers and CBM10-1 to CBM10-3: 323-689 and GH6: amino acids 690-1103), the colors of the lines indicating sequence identity to query. The lower panels display the predicted Local Distance Difference Test (LDDT) of the individual amino acids in the five predicted models (the model ranked "1", was used in this study).

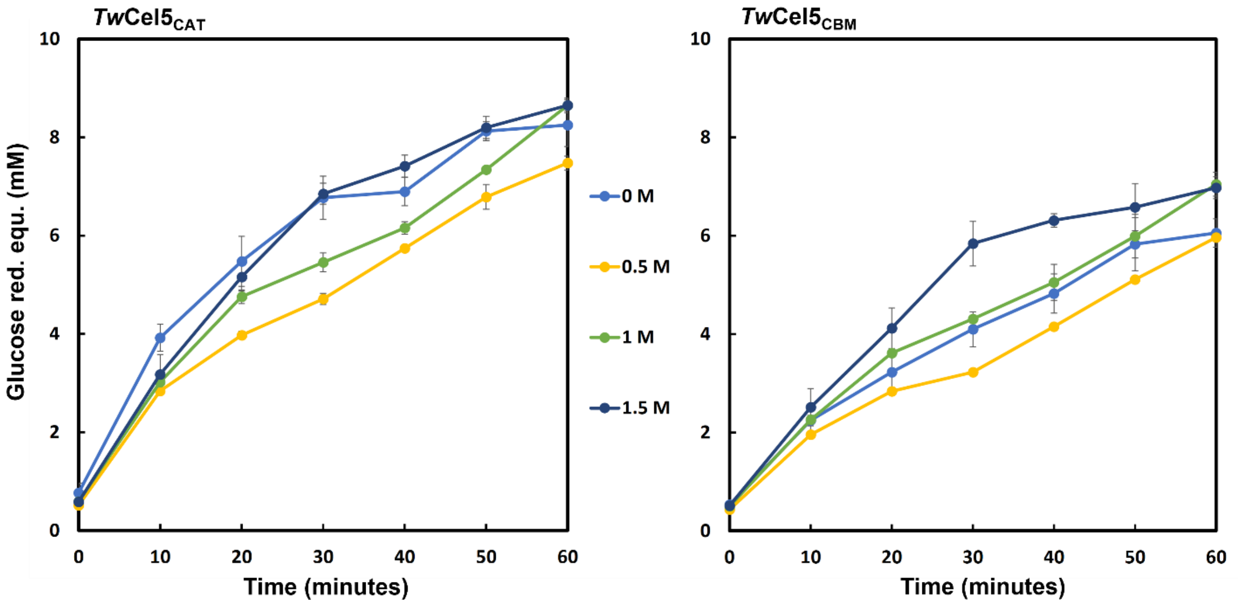


**Fig. S2.** Effect of NaCl concentration on hydrolytic activity of *Tw*Cel5$_{CAT}$ and *Tw*Cel5$_{CBM}$ measured using β-glucan (0.5% w/v) in 50 mM potassium phosphate buffer at pH 7.5, 50 nM enzymes and varying concentration of NaCl (0 - 1.5 M) incubated at 30º for 60 minutes. Reducing sugar equivalents were calculated by DNSA assay using glucose as standard. Values shown are mean and standard deviation obtained from triplicates.

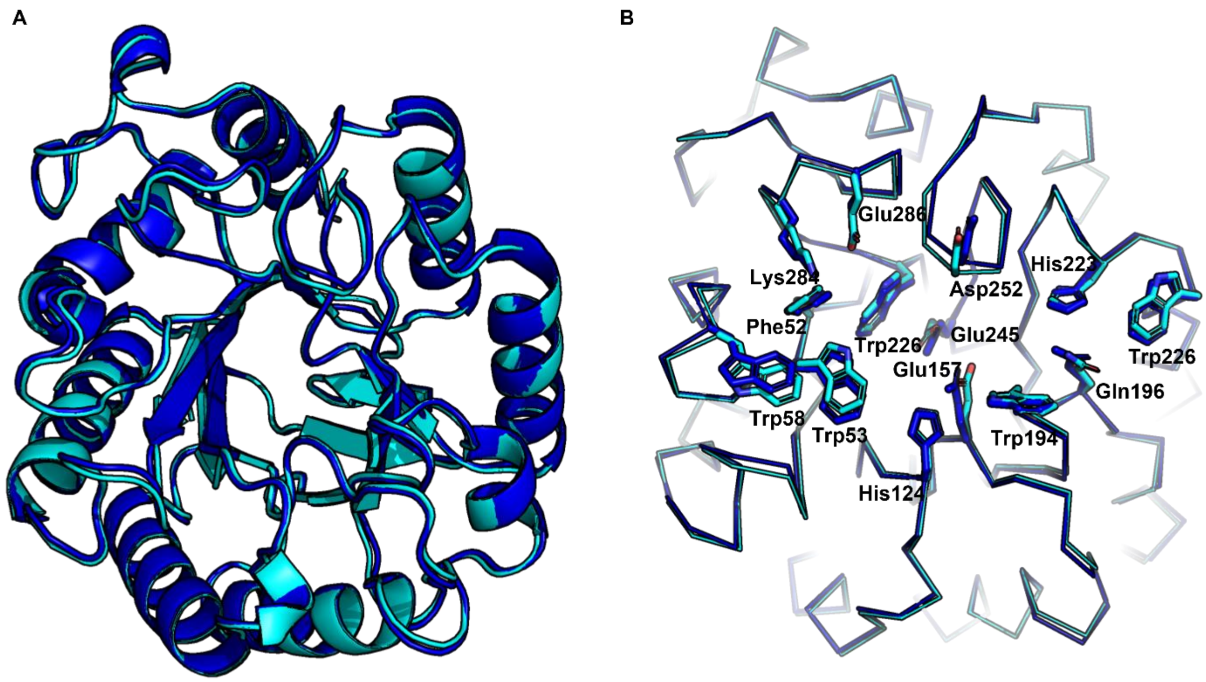


**Fig. S3.** Comparison of the experimentally determined structure of *Tw*Cel5$_{CAT}$ and its Alphafold2 model. (A) Cartoon representation of the X-ray crystallographic model (cyan) and the Alphafold2 predicted model (blue) structurally superimposed. (B) Comparison of conserved amino acids in the active site and substrate binding cleft.

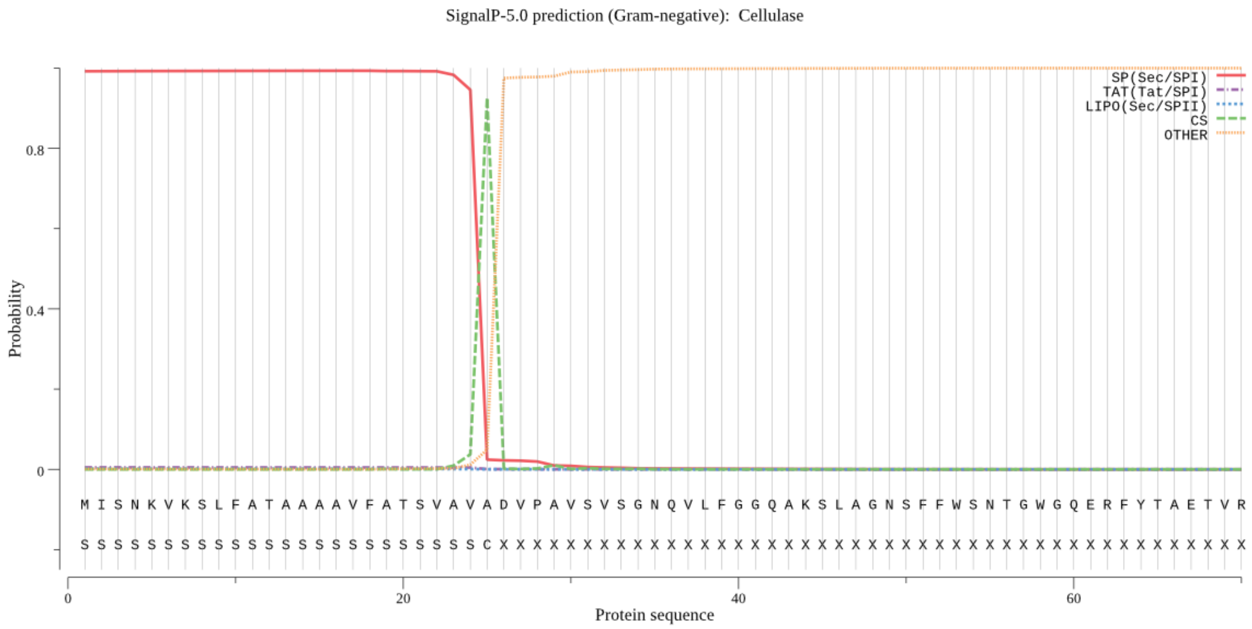


**Fig. S4.** Prediction of the putative signal peptide in deduced amino acid sequence of *Tw*Cel5-6 (residues 1-25) based on the SignalP-5.0 prediction tool.

**Table S1.** Primers used for amplification of gene encoding the *Tw*Cel5 variants.

| Gene | Primers | Primer sequence (5'-3') | Vector name |
|---|---|---|---|
| $TwCel5_{CAT}$ | TwCel5CATf<br><br>TwCel5CATr | TTAAGAAGGAGATATACTATGGCGTTTAGCCAGGGTGC<br><br>AATGGTGGTGATGATGGTGCGCCATACCCCACAGCAGTTTCA | pNIC-CH |
| $TwCel5_{CBM}$ | TwCel5CBMf<br><br>TwCel5CBMr | CACCGTGAGTGTTTCCGGTAATCA<br><br>TTAGGTACAGCTCGAGATAACACCG | pET151/D-TOPO |
